# Prokaryotic virus Host Predictor: a Gaussian model for host prediction of prokaryotic viruses in metagenomics

**DOI:** 10.1101/2020.12.02.408310

**Authors:** Congyu Lu, Zheng Zhang, Zena Cai, Zhaozhong Zhu, Ye Qiu, Aiping Wu, Taijiao Jiang, Heping Zheng, Yousong Peng

## Abstract

**Background:** Viruses are ubiquitous biological entities, estimated to be the largest reservoirs of unexplored genetic diversity on Earth. Full functional characterization and annotation of newly-discovered viruses requires tools to enable taxonomic assignment, the range of hosts, and biological properties of the virus. Here we focus on prokaryotic viruses, which include phages and archaeal viruses, and for which identifying the viral host is an essential step in characterizing the virus, as the virus relies on the host for survival. Currently, the method for determining the viral host is either to culture the virus, which is low-throughput, time-consuming, and expensive, or to computationally predict the viral hosts, which needs improvements at both accuracy and usability. Here we develop a Gaussian model to predict hosts for prokaryotic viruses with better performances than previous computational methods.

**Results:** We present here Prokaryotic virus Host Predictor (PHP), a software tool using a Gaussian model, to predict hosts for prokaryotic viruses using the differences of *Æ*-mer frequencies between viral and host genomic sequences as features. PHP gave a host prediction accuracy of 34% (genus level) on the VirHostMatcher benchmark dataset and a host prediction accuracy of 35% (genus level) on a new dataset containing 671 viruses and 60,105 prokaryotic genomes. The prediction accuracy exceeded that of two alignment-free methods (VirHostMatcher and WIsH, 28%-34%, genus level). PHP also outperformed these two alignment-free methods much (24%-38%*vs* 18%-20%, genus level) when predicting hosts for prokaryotic viruses which cannot be predicted by the BLAST-based or the CRISPR-spacer-based methods alone. Requiring a minimal score for making predictions (thresholding) and taking the consensus of the top 30 predictions further improved the host prediction accuracy of PHP.

**Conclusions:** The Prokaryotic virus Host Predictor software tool provides an intuitive and user-friendly API for the Gaussian model described herein. This work will facilitate the rapid identification of hosts for newly-identified prokaryotic viruses in metagenomic studies.

**Author Summary:** Prokaryotic viruses which include phages and archaeal viruses play an important role in balancing the global ecosystem by regulating the composition of bacteria and archaea in water and soil. Identifying the viral host is essential for characterizing the virus, as the virus relies on the host for survival. Currently, the method for determining the viral host is either to culture the virus which is low-throughput, time-consuming, and expensive, or to computationally predict the viral hosts which needs improvements at both accuracy and usability. This study developed a Gaussian model to predict hosts for prokaryotic viruses with better performances than previous computational methods. It will contribute to the rapid identification of hosts for prokaryotic viruses in metagenomic studies, and will extend our knowledge of virus-host interactions.

## Background

Viruses are ubiquitous biological entities, with an estimate of 10^31^ viral particles at any given time on earth [1]. They infect all types of organisms, from animals and plants to bacteria and archaea. Prokaryotic viruses which include phages and archaeal viruses play an important role in balancing the global ecosystem by regulating the composition of bacteria and archaea in water and soil [2–5]. For humans, phages directly influence gut health and are associated with several human diseases, such as diabetes [6] and Crohn’s disease [7]. Interestingly, phages can also be applied as therapy for bacterial infections [8, 9], especially for bacterial strains resistant to multiple antibiotics.

Viruses are estimated to be the largest reservoirs of unexplored genetic diversity on earth [5]. Novel viruses have been discovered at an unprecedented pace with the rapid expansion of metagenomics [4, 10–12]. For example, the recent Tara Oceans Project greatly extended the global ocean DNA virome by identifying 195,728 viral populations [4]. As our knowledge of the viral genetic diversity expands considerably, it becomes increasingly critical to develop tools to facilitate functional characterization and annotation of the newly-discovered viruses, such as taxonomic assignment, range of hosts, biological properties, and so on.

Identifying the viral host is essential for characterizing the virus, as the virus relies on the host for survival. Currently, the method for determining the viral host is to culture the virus, which is low-throughput, time-consuming, and expensive [13]. Even worse, few viruses can be cultured since less than one percent of microbial hosts have been cultivated successfully in laboratories [14, 15]. Therefore, it is highly desirable to develop quicker methods for predicting hosts of newly discovered viruses in metagenomic studies.

Several computational approaches have been introduced to predict hosts of viruses based on viral genomic sequences. They can be largely classified into two groups according to their dependence on alignment: alignment-based methods and alignment-free methods. The alignment-based methods rely on sequence similarity searches between a query virus and candidate host genomes because viruses and their hosts sometimes share genes and/or short nucleotide sequences [16, 17]. Such sequences may come from spacer sequences used in CRISPR systems, integration sites used by proviruses or horizontal gene transfer. BLAST is most widely used to predict viral hosts with relatively high accuracy [16, 17] based on the similarity between virus and host genomes. However, for newly identified viruses divergent from the known ones, the applicability of the BLAST-based method could be limited. The CRISPR-spacer-based method has shown a higher accuracy compared to the BLAST-based method in predicting phage hosts, yet it can only be used for a small proportion of viruses since only 40%-70% prokaryotes encode a CRISPR system [17] and not all of them have spacer sequences from viruses. Besides, since the spacers are only the infection history of an individual prokaryotic cell, a precise phage-bacteria sequence match would require the unlikely preservation of the CRISPR-spacers.

The alignment-free methods predict the host of viruses based on co-occurring *k*-mers to other phages with known hosts [18], or sequence composition similarity between viruses and their hosts. Among the latter kind of methods, VirHostMatcher (VHM) [16] and WIsH [19] have achieved the highest host prediction accuracy. VHM employs the background-subtracting measure d_2_^*^ for measuring the similarity of oligonucleotide frequency between viruses and hosts, and predicts the one with the lowest distance to the virus as the viral host. VHM achieved a host prediction accuracy of 33% at the genus level. WIsH predicts viral hosts by training a homogeneous Markov model for each potential host genome, then calculating the likelihood of a contig under each of the trained Markov models, and finally predicting the host whose model yields the highest likelihood. WIsH has achieved 63% mean accuracy at the genus level on their benchmark dataset of 3kbp phage contigs, and is currently the best tool for predicting phage hosts based on short phage contigs.

In this study, a Gaussian model (GM) for predicting hosts of prokaryotic viruses was developed based on the differences of *k*-mer frequencies between viral and host genomic sequences. A standalone tool and a web-based tool was implemented to run the GM. The GM not only outperformed previous alignment-free methods, but also shaped a complement to the alignment-based methods in predicting hosts for prokaryotic viruses. The GM should facilitate the prediction of virus host in metagenomic studies.

## Methods

### Datasets of virus-host interactions

The prokaryotic viruses were referred to as viruses hereinafter unless otherwise specified. Two datasets were used to build and test computational models for predicting virus hosts. The first was the VirHostMatcher (VHM) benchmark dataset obtained from Ahlgren’s study [16]. The taxonomic information of both viral and prokaryotic genomes in the dataset was updated according to the International Committee on Taxonomy of Viruses (ICTV) [20] and NCBI Taxonomy database [21]. One pair of virus-host interaction (Tetraselmis viridis virus - Tetraselmis sp.) was removed due to the incorrect annotation of the Tetraselmis as bacteria. The updated VHM dataset contains a total of 1,426 pairs of virus-host interactions, 1,426 virus genomes, and 31,918 prokaryotic genomes. It was available to the public on GitHub [22].

The second dataset was the test dataset to assess the computational models of predicting virus hosts. Contrary to the VHM dataset that contains virus-host interactions compiled from the NCBI RefSeq database [23] before May 5^th^, 2015, the test dataset contains those submitted between May 6^th^, 2015 and February 26^th^, 2019. The virus-host interactions which have both the same viral species and the same host genus with those in the VHM dataset were removed. The test dataset contains a total of 671 pairs of virus-host interactions, 671 virus genomes, and 60,105 prokaryotic genomes obtained from the NCBI genome database [24] on February 21^th^, 2019. The taxonomy distribution of both the virus and host in the test dataset and the VHM dataset was analyzed and shown in Additional file 1: Figure S1. When compared to the VHM dataset, the test dataset includes 667, 97 and 2 new viral species, genus and families, respectively, and 37, 11 and 8 new host species, genus and families, respectively.

### The Gaussian model for predicting virus host

The Gaussian mixture model is a probabilistic model which uses a finite number of Gaussian distributions to fit data points and get the probability density of them [25]. Here, the Gaussian mixture model with only one component was found to perform best in predicting virus hosts (Additional file 1: Figure S2); therefore, the Gaussian mixture model was simplified as Gaussian model (GM). The GM takes the differences of *k*-mer frequencies between virus and prokaryotic genomic sequences as features, and outputs a score (the logarithm of the probability of being viral host) for the prokaryote. The *k*-mers of 4 nucleotides were selected (Additional file 1: Figure S2), which resulted in 256 features. The GM was built using the function of GaussianMixture in scikit-learn [25, 26]

### Definition of accuracy in virus host prediction by Gaussian models

For each virus, the GM calculated a score (the logarithm of the probability of being viral host) for all prokaryotic genomes available in the dataset. For example, in the test dataset, each of the 60,105 prokaryotic genomes would be assigned a score (the logarithm of the probability of being viral host) by the GM. The prokaryotic species with the highest score was considered as the predicted host of the virus. The predicted host was compared to the actual host at different taxonomic levels. If the predicted host belonged to the same taxonomic unit such as genus with the actual host, the prediction was considered as correct at the level. The accuracy of virus host prediction at a certain taxonomic level was defined as the ratio of correctly predicted host at this taxonomic level.

### Prediction of virus hosts with VHM and WIsH

VHM and WIsH were the best alignment-free methods for predicting phage hosts according to previous studies [16, 19]. For comparison, they were tested with default parameters on the test dataset mentioned above. They were computed with the codes available at GitHub [27, 28].

### Prediction of virus hosts with alignment-based methods

Previous studies by Edwards et al. [17] showed the sequence alignment-based methods, such as the BLAST-based method and CRISPR-spacer-based methods, achieved excellent performances in predicting virus hosts. We compared these methods with the GM in predicting virus hosts on the test dataset.

To predict the virus host based on BLAST, the genome sequence of each virus was queried against the prokaryotic genomes in the test dataset using blastn (version 2.6.0+) [29]. The prokaryotic species with the best hit which had E-value smaller than 1E-5 was considered as the potential host of the virus.

To predict the virus host based on CRISPR spacer sequences, firstly, the CRISPR spacer sequences in all prokaryotes genomes of the test dataset were extracted by the CRISPR Recognition Tool (CRT) [30]. Then, the genome sequence of each virus was queried against the prokaryotic CRISPR spacer sequences using blastn (version 2.6.0+). The hits, i.e., the CRISPR spacer sequences, with identity > 95% to the query sequence over the whole spacer length were considered as perfect hits. The prokaryotic species with perfect hits to the virus genome was considered as the potential host of the virus.

### Prediction of virus hosts based on simulated viral contigs

Metagenomic assembly often yields partial genomes, so the prediction of the virus host based on contigs of varying lengths was important. To test the GM in prediction of virus hosts in metagenomics, the GMs based on simulated viral contigs of length L bp (L =1,000, 3,000, 5,000 and 10,000) randomly subsampled from viral genomes were built, and the models were evaluated by the ten-fold cross-validations on the K-means clustering of the VHM dataset.

## Results

### Building a Gaussian model for predicting virus hosts

Viruses and their hosts often share similar oligonucleotide frequency patterns in their genomes, yet the prediction of virus-host interactions based on the similarity pattern remains challenging. A Gaussian model (GM) was developed in this study to predict the hosts based on the differences of *k*-mer frequencies between viral and host genomic sequences (see Methods). To evaluate the ability of the GM in predicting virus hosts, a strict testing strategy was adapted (Figure 1). Firstly, a feature vector characterizing the differences of *k*-mer frequencies between viral and host genomic sequences was calculated for each pair of virus-host interaction within the VHM dataset. The K-means algorithm was then used to separate the virus-host interactions in the VHM dataset into ten clusters based on the feature vectors. Finally, ten-fold cross-validations were conducted as follows: nine clusters of virus-host interactions were used to train the GM; while the outcome GM model was then used to predict the virus-host interactions in the remaining cluster. For each virus, scores were assigned to all the prokaryotic host species in the VHM dataset, and the prokaryotic species with the highest score was predicted to be the host of the virus. The above process was repeated for each cluster. The overall prediction accuracy of the GM was calculated as the ratio of the correctly predicted viruses among all viruses in the dataset.

**Figure 1.**
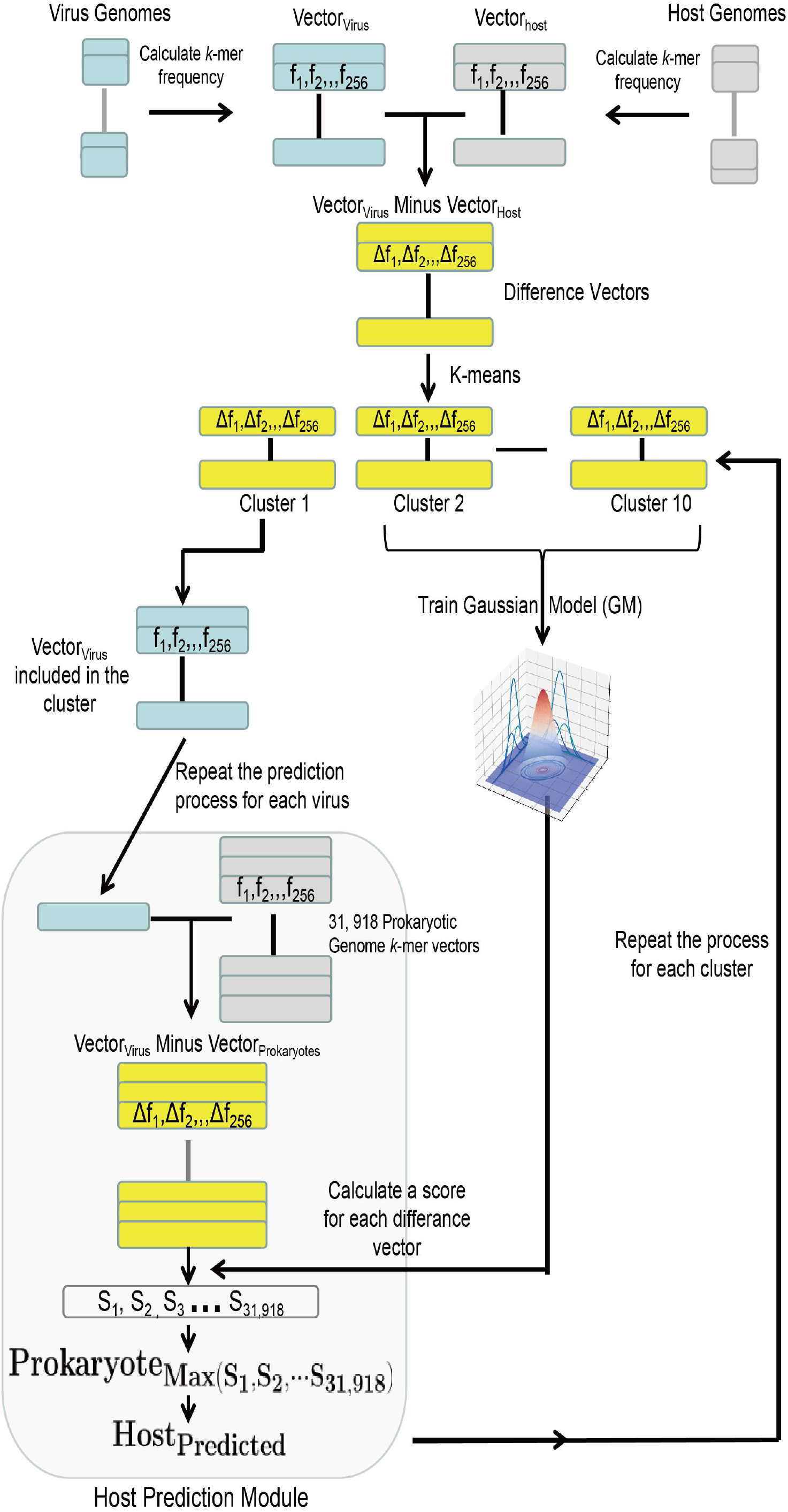
The testing strategy for the GM on the VHM dataset.

The testing strategy mentioned above was also used to determine parameter values for the GM. Two important parameters for the GM was the length of *k*-mers and the number of components (i.e., the number of Gaussian distribution) used in the model. The virus host prediction accuracy of the GM increased as the length of *k*-mers increased from 1 to 5, and then decreased with *k*-mers of six nucleotides (Additional file 1: Figure S2A). The *k*-mers with four nucleotides which have a total of 256 kinds of *k*-mers were selected to balance the model complexity and prediction accuracy since the number of samples used in training the GM is only 1,426. When selecting the number of components used in the GM, interestingly, we found the GM with one component outperformed that with multi-components (Additional file 1: Figure S2B). Therefore, the GM with one component and with *k*-mers of four nucleotides was used in the further analysis. The optimized GM had a virus host prediction accuracy of 0.34 on the genus level and 0.45 on the family level in the ten-fold cross-validations on the K-means clustering of the VHM dataset (Figure 2). For comparison, the prediction accuracies of VHM and WIsH on the VHM dataset were also displayed. The GM slightly outperformed VHM and WIsH on all taxonomic levels (Figure 2).

**Figure 2.**
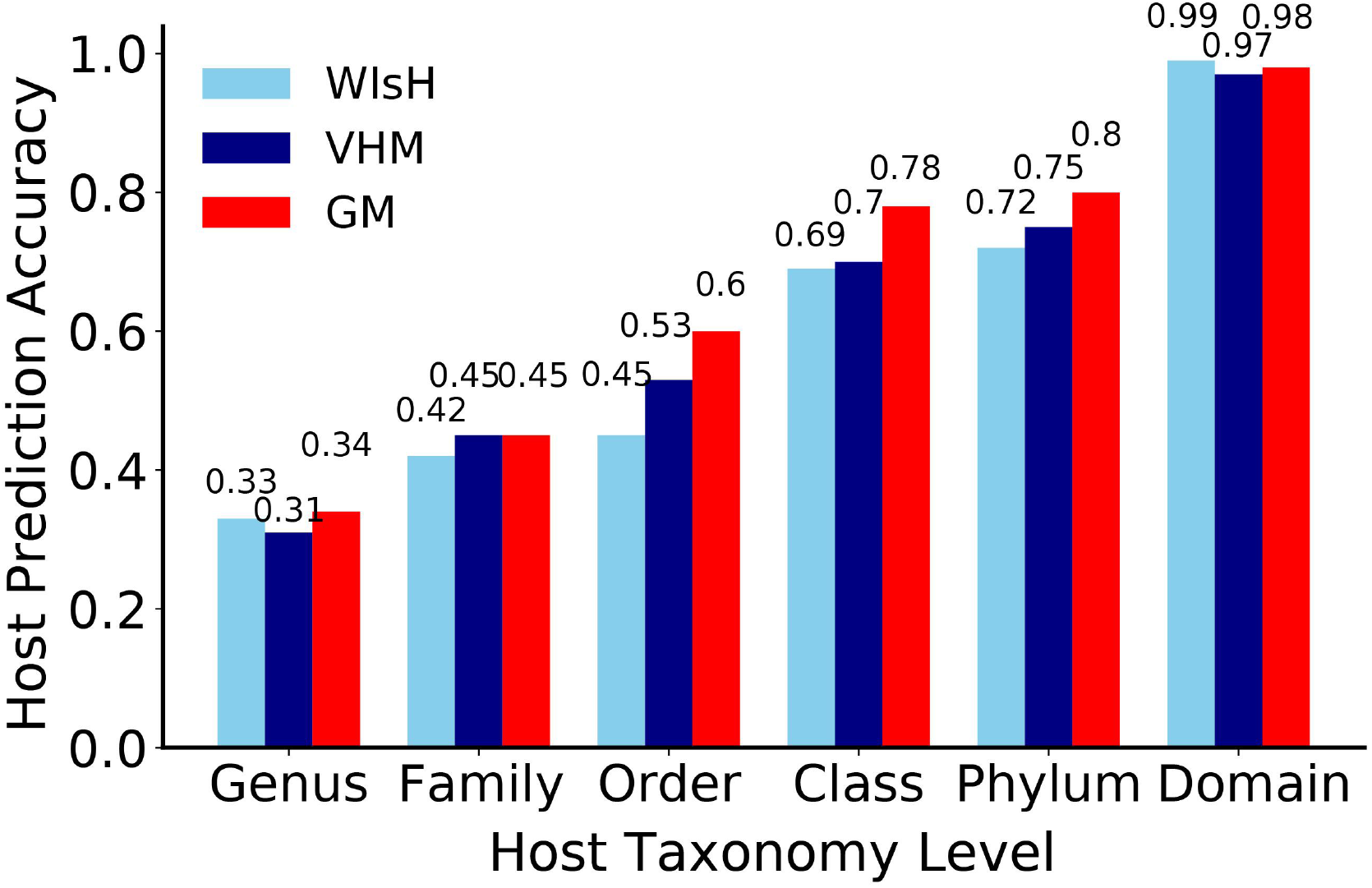
The virus host prediction accuracies of the GM in the ten-fold cross-validations on the K-means clustering of the VHM dataset, and its comparison to VHM and WIsH.

The GM was also compared to other common machine-learning algorithms in predicting virus hosts, including the random forest, logistic regression, naive Bayesian, decision tree, k-nearest neighbor and multi-layer perceptron algorithms. The GM was found to outperform much than these machine-learning algorithms in the ten-fold cross-validations on K-means clustering of the VHM dataset (Additional file 1: Figure S3).

### Prediction performances of the GM on the test dataset

The GM built on the VHM dataset was further tested using the test dataset. The predictive accuracy of the GM increased with the gradual elevation of the taxonomic level from genus to phylum (Figure 3A). Notably, the GM achieved accuracies of 0.46 on the level of genus and 0.63 on the level of family. For comparison, VHM [16] and WIsH [19] were also used to predict the prokaryotes host on the same test dataset. Our GM achieved much higher accuracies than these two methods at all taxonomic levels. For example, the prediction accuracies of GM at the genus level was 0.18 and 0.12 higher than VHM and WIsH, respectively (Figure 3A).

**Figure 3.**
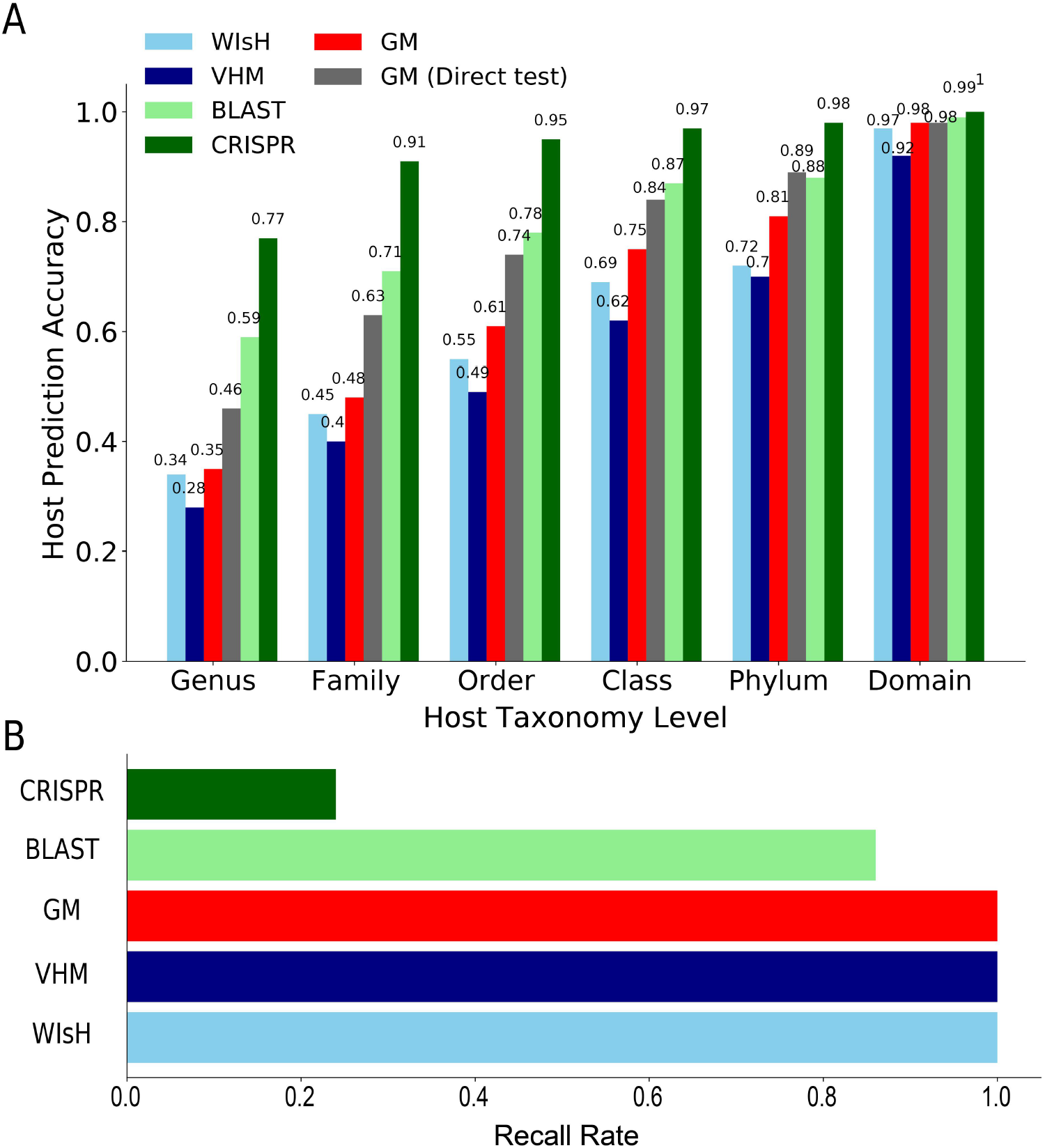
The host prediction accuracy (A) and the recall rate (B) of the GM model and their comparisons to other computational methods on the test dataset.

The shared viruses and hosts in both the test dataset and the training dataset (VHM dataset) may result in over-estimate of the performance of the GM in the test dataset. Therefore, when predicting a target virus-host interaction in the test dataset, the GM was re-built based on the training dataset from which the virus-host interactions that shared the same viral and host genus with the target virus-host interactions were removed. This resulted in a compromised performance of the GM on the test dataset (see red bars in Figure 3A). For example, the predictive accuracies of the GM were 0.35 on the genus level and 0.48 on the family level. However, most of the time, the GM still outperformed both VHM and WIsH (Figure 3A).

The similarity between the virus and host genomic sequences often indicates virus-host relations. Thus the alignment-based methods, such as the BLAST-based method and the CRISPR-spacer-based method, are frequently used to predict the virus host. These two methods were tested on the test dataset and were compared to the GM (Figure 3). The CRISPR-spacer-based method predicted virus hosts with the highest accuracies at all taxonomic levels, ranging from 0.77 to 1, among all methods, but it can only predict hosts for less than one-fourth of viruses. The BLAST-based method predicted hosts for most viruses with accuracies higher (1%-13%) than those of the GM.

We further investigated the performance of the alignment-free methods in predicting hosts for viruses which cannot be predicted by the alignment-based methods on the test dataset. A total of 48 viruses cannot be predicted by the BLAST-based method. The GM predicted hosts more accurately than both VHM and WIsH at all taxonomic levels for these viruses (Figure 4A). For example, the GM had a prediction accuracy of 0.24 at the genus level, while the VHM and WIsH had accuracies of 0.18 and 0.20, respectively. A total of 430 viruses cannot be predicted by the CRISPR-spacer-based method. The GM again had much higher accuracies than both VHM and WIsH at all taxonomic levels on these viruses. These results suggest that the GM may be a better complement to the alignment-based methods than the VHM and WIsH.

**Figure 4.**
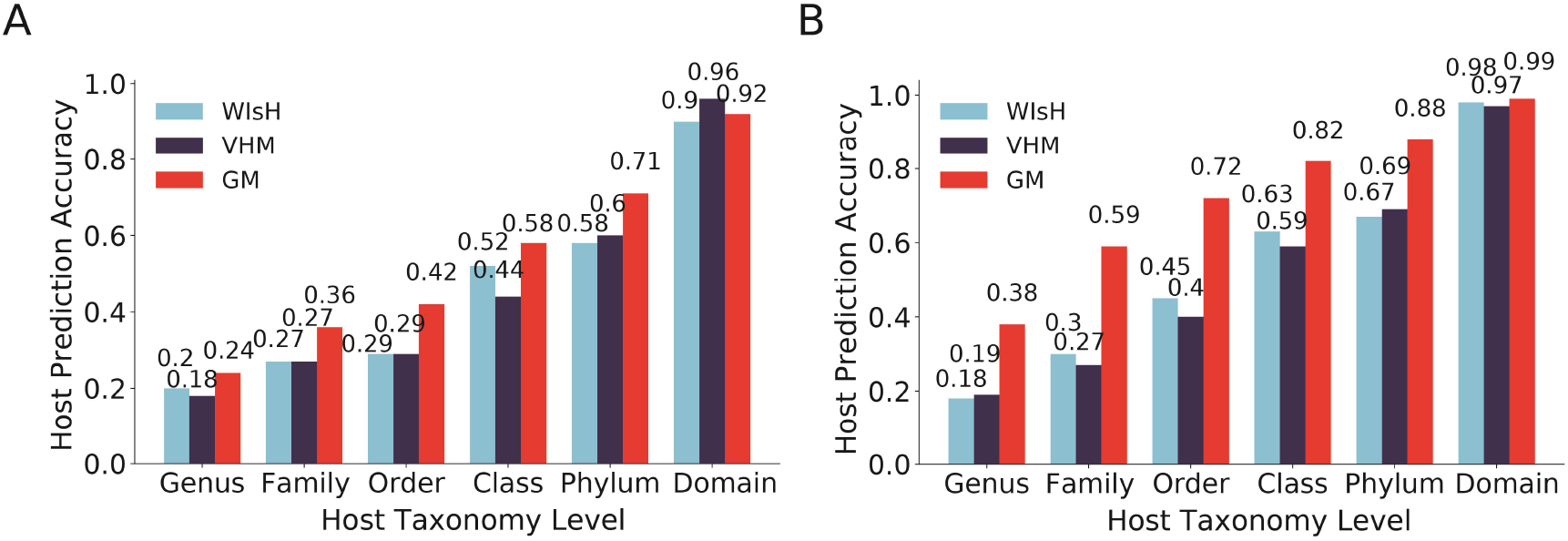
The performance of the alignment-free methods in predicting hosts for viruses which cannot be predicted by the BLAST-based (A) and the CRISPR-spacer-based methods (B).

### Approaches for further improvements of the GM

To further improve the performance of GM in host prediction, the maximum consensus method was applied as in Ahlgren’s work [16]. The predicted host taxon was selected as the most frequent taxon among the N (N = 1 to 30) hosts with the highest score. When 30 predicted hosts were considered, the prediction accuracy at the genus level improved significantly using this consensus approach for GM and VHM, achieving an accuracy of 0.45 and 0.39, respectively (Figure 5A & Additional file 1: Table S1); while the prediction accuracy of WIsH at the genus level increased as N increased from 1 to 10, then it began to decrease. Similar variation patterns were observed at other taxonomic levels for all three alignment-free methods (Additional file 1: Table S1).

**Figure 5.**
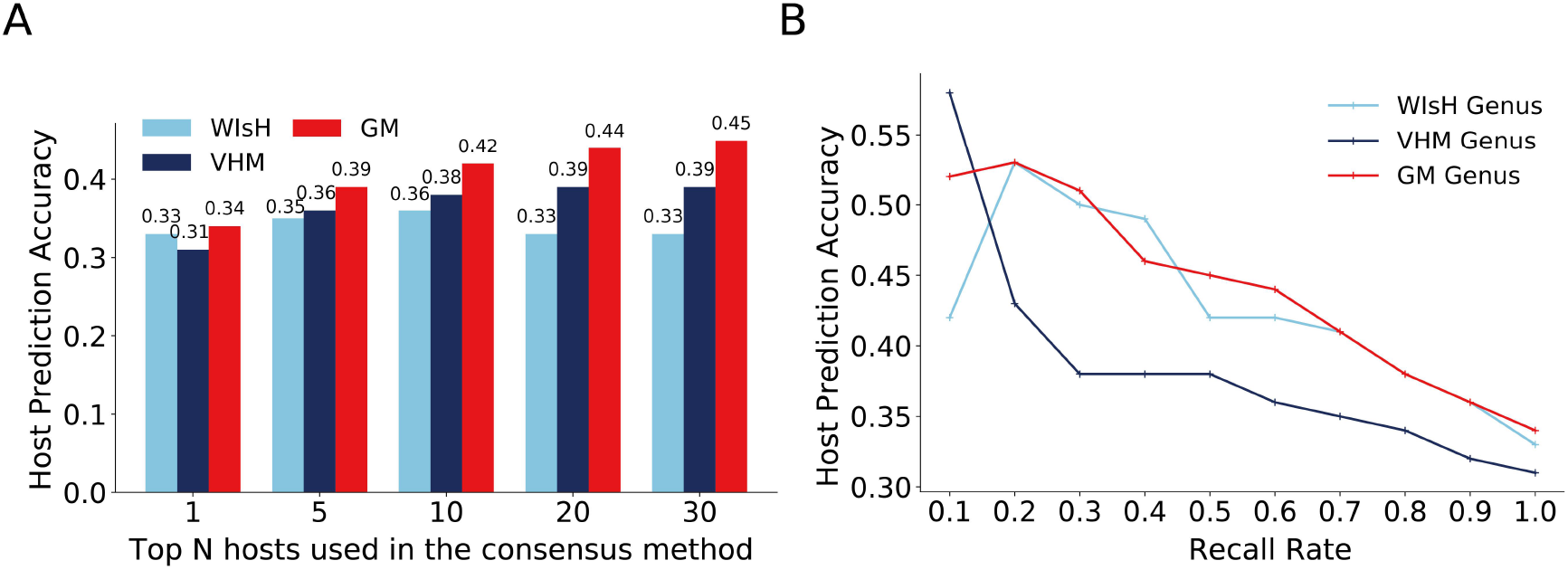
Improvement of the GM by the consensus method (A) and threshold method (B) on the VHM dataset. The host prediction accuracies of the GM was obtained from the ten-fold cross-validations on the K-means clustering of the VHM dataset. Only the prediction accuracies at the genus level were shown in the figure for all methods. The host prediction accuracies at higher taxonomic levels were shown in Additional file 1: Table S1 and S2.

**Figure 6.**
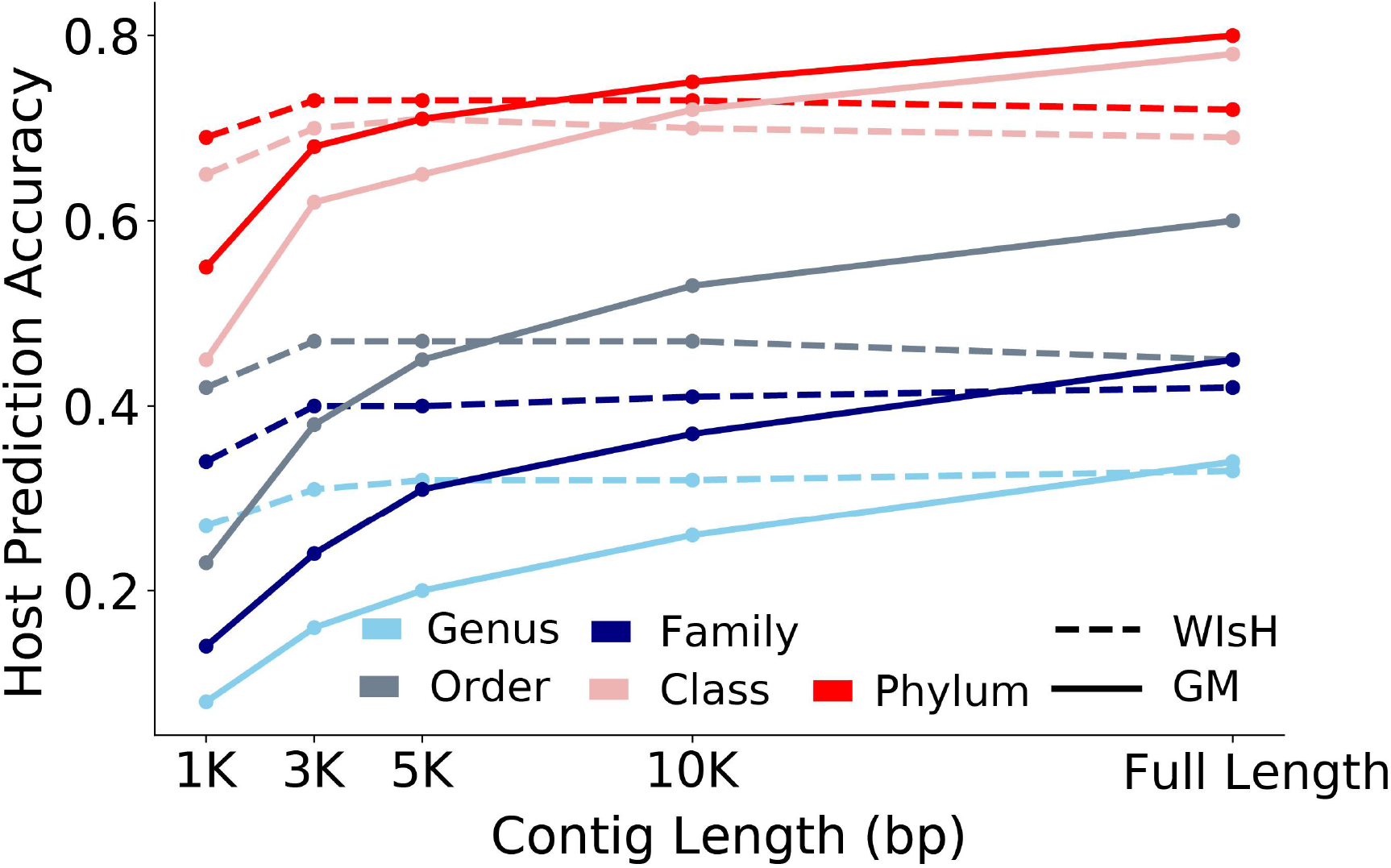
Host prediction accuracy of GM (solid line) and WIsH (dashed line) at all taxonomic levels based on viral contigs of varying lengths. The host prediction accuracies of the GM were obtained in the ten-fold cross-validations on the K-means clustering of the VHM dataset.

We also tested applying a score threshold requirement to making host predictions as Ahlgren et al. did in their study [16]. Predictions were only made when the score was larger than a given threshold. The host prediction accuracies and the recall rate of the GM, VHM and WIsH were calculated (Figure 5B & Additional file 1: Table S2). As was shown in Figure 5B, the prediction accuracy of the GM, VHM and WIsH at the genus level increased significantly as the recall rate decreased. Both GM and WIsH outperformed VHM much as the recall rate ranging from 1 to 0.2. Interestingly, VHM outperformed much than GM and WIsH at the recall rate of 0.1. The GM slightly outperformed WIsH when the recall rate ranged from 1 to 0.5. Further analysis of the prediction accuracies at higher taxonomic levels versus the recall rate found that the GM outperformed VHM and WIsH much at both the family and order level when the recall rate ranged from 1 to 0.1; while at other taxonomic levels, these methods performed comparably (Additional file 1: Table S2).

### Host prediction based on viral contigs

Metagenomic assembly often yields partial genomes. By far WIsH was reported as the most accurate method for predicting phage hosts based on short contigs [19]. We further tested the ability of the GM in predicting hosts based on viral contigs of varying length ranging from 1kb to 10kb. When testing the GM and WIsH on the VHM dataset, we found that the GMs achieved lower accuracies than WIsH at all taxonomic levels when viral contigs were smaller than 10kb; when viral contigs were equal to or larger than 10kb, the GMs had higher accuracies than WIsH at order or higher taxonomic levels.

### Development and application of a software tool for predicting host of prokaryotic viruses based on the GM

We developed a software tool named Prokaryotic virus Host Predictor (PHP) to provide a friendly user interface for the GM algorithm. PHP is freely available either as a standalone version [22] or in the form of a web server [31]. We included both the VHM and the test dataset for GM model training in PHP to maximize the usability based on all currently available data. While the standalone version of PHP is suitable for host prediction of a large number of viruses, the PHP web server is suitable for host prediction of fewer than 100 viruses. The web server of PHP is intuitive and user-friendly. It takes one or multiple virus genomic sequences as input. After submission, a waiting page appears and would last from several minutes to several hours depending on the number and size of viral genomic sequences. The user can bookmark the page and check the status of the job in the “Job status” page, or provide the email address (optional) and check the results upon email notification. The output would show the name, the score, and the taxa (from species to phylum) of the predicted host for the given viruses. Both the top 1 and the consensus of top 5 predicted hosts were shown since considering the consensus of the top 5 predictions would improve the performances of the GM.

The time consumed by the PHP was measured on the VHM dataset, and was compared to the time consumed by WIsH which was reported to predict phage host rapidly. When tested on a laptop with 8 threads (see Additional file 1: Table S3 for details), the process of model building of PHP took 3 hours 30 minutes, which included the calculation of *k*-mer frequencies in viral and prokaryotic genomes, and model training, while the process of model building of WIsH took 27 minutes (Additional file 1: Table S3); However, the PHP only used half of the time (57 minutes) in host prediction for 1,426 viruses when compared to WIsH (1 hour and 51 minutes). Overall, the total time consumption of PHP was double to that of WIsH (Additional file 1: Table S3). When tested on a server with 40 threads (see Additional file 1: Table S3 for details), the time consumption of both PHP and WIsH was reduced much compared to that of the tests on the laptop. For example, the time consumption of PHP was reduced from 3 hours 30 minutes to 48 minutes during the process of model building, while that of WIsH was reduced from 27 minutes to 12 minutes (Additional file 1: Table S3). However, the reduction of time consumption of PHP was larger than that of WIsH, and the total time consumption of PHP was less than that of WIsH (1 hours 1 minutes *vs* 1 hours 14 minutes).

Finally, we tested the ability of the PHP in predicting virus hosts using 139 pairs of known phage-host interactions which were determined by the single-cell viral tagging method [32]. These pairs of phage-host interactions were available at GitHub [22]. The online version of the PHP predicted hosts for these phages with an accuracy of 0.29 on the genus level and 0.64 on the family level (Table 1). When considering the top 5 predictions, the prediction accuracy increased to 0.33 on the genus level and 0.75 on the family level (Table 1). Considering that the bacterial contigs identified in the same project with the viral contigs are more likely to be the host of these viruses, the local version of the PHP was used to predict hosts for these phages among the 289 bacterial contigs used in the same study. The prediction accuracies were further improved to 0.67 on the genus level and 0.80 on the family level (Table 1).

**Table 1.** The prediction accuracies of the PHP in predicting phage hosts for 139 pairs of known phage-host interactions obtained by the single-cell viral tagging method.

| PHP with different usage mode | Genus | Family | Order | Class | Phylum |
| --- | --- | --- | --- | --- | --- |
| Online version (top 1) | 0.29 | 0.64 | 0.85 | 0.85 | 0.94 |
| Online version (top 5) | 0.33 | 0.75 | 0.83 | 0.83 | 0.93 |
| Local version, with 289 bacterial<br>contigs | 0.67 | 0.80 | 0.84 | 0.85 | 0.86 |

## Discussion

In this study, we developed the GM to predict the viral hosts based on the difference of *k*-mer frequencies between virus and host genomes. On the genome-scale, the GM performed better than VHM and WIsH on both the VHM benchmark dataset and the test dataset. Although the GM had lower host prediction accuracies than the alignment-based methods, it can predict hosts for all prokaryotic viruses. Besides, for those viruses which cannot be predicted by the alignment-based methods, the GM outperformed VHM and WIsH much, suggesting that the GM can be a more suitable complement to the alignment-based methods than VHM and WIsH. The GM can be further improved by the consensus and threshold methods. These results suggest that the GM are useful for host prediction of viruses in metagenomic studies.

Viruses and their host genomes often share similar oligonucleotide frequency patterns, but it is challenging to select the best metric for measuring the similarity. Several common metrics have been used for measuring the similarity of oligonucleotide frequencies between viral and host genomes, such as the Euclidean distance and Manhattan distance [16]. Previous studies by Ahlgren et al. [16] comprehensively compared 11 common oligonucleotide frequency metrics for predicting viral hosts, and found the background-subtracting measure d_2_^*^ performed best among these metrics. Compared to previous studies, this study is unique in that the GM developed herein learned a best “metrics” to measure the similarity between viral and host genomes. The GM took the differences of *k*-mer frequencies between viral and host genomic sequences as features in viral hosts prediction. The GM does not detect patterns specific to a genome or a pair of genomes, but rather detect patterns in the difference of the *k*-mer frequencies between a virus and a host genome. The GM with only one Gaussian distribution has the highest performance, suggesting that the patterns in the difference of the *k*-mer frequencies between virus and host genomes exhibit similar features. GM outperformed both VHM and WIsH in viral host prediction, indicating that it can learn a more suitable “metrics” for measuring the similarity between *k*-mer frequencies of viral and host genomes than the existing metrics.

Multiple common machine-learning algorithms were used to predict virus hosts based on the differences of *k*-mer frequencies between virus and host genomic sequences. However, the GM outperformed much than other algorithms (Additional file 1: Figure S3). The possible reason is that the differences between the *k*-mer frequencies of virus and host genomes are supposed to be close to zero since viruses and their hosts often share similar oligonucleotide frequency patterns in their genomes. The *k*-mer frequency differences between virus and host genomes approximately followed a normal distribution with a mean of zero, and were different from those between virus and non-host prokaryotic genomes (Additional file 1: Figure S4). Therefore, the GM is more suitable for capturing the *k*-mer frequencies differences between virus and host genomes than other machine-learning algorithms.

Accurate prediction of virus hosts is challenging. Lots of computational methods have been developed for prediction of virus hosts in previous studies. Among the methods tested in this study, the CRISPR-spacer-based method is the most accurate one, but it only predicted host for less than one-fourth of viruses. The BLAST-based method can predict host for most viruses, and outperformed the alignment-free methods including GM, VHM and WIsH. It should be a promising tool for predicting virus hosts considering the easy use of BLAST. The alignment-free methods have the advantage of a high recall rate. Both VHM [16] and WIsH [19] are independent of training and are supposed to be more robust in applications, while the GM needs training and may have the risk of over-fitting. To reduce over-fitting of the GM, a strict testing strategy of ten-fold cross-validations on the K-means clustering was used to evaluate the performance of GM on the VHM benchmark dataset (Figure 1). The GM outperformed existing alignment-free methods (VHM and WIsH) on both the benchmark dataset and the test dataset (Figure 2&3), especially for those viruses which cannot be predicted by the alignment-based methods (Figure 4). Taken together, a combination of multiple methods, including the alignment-free methods and the alignment-based methods, would help further improve the prediction of virus hosts.

A major bottleneck of this study is the limitation and bias of the dataset of virus-host interactions, considering the huge diversity of prokaryotic viruses on the earth [4, 10]. The virus-host interactions in our datasets are biased towards some common viruses and host taxa, which reflects the taxonomic distribution of viruses and prokaryote in nature. For example, the three most commonly observed families (Siphoviridae, Myoviridae, and Podoviridae) account for 77% of all viruses in the VHM dataset (Additional file 1: Figure S5) and are indeed the most commonly observed phage taxa in nature [4, 33]. Another limitation of the study is that the GM was inferior to WIsH in predicting virus hosts on the viral contigs less than 10kb. Importantly, the GM has a great potential of improvements when given a more diverse and high-quality training dataset which would be enabled by more effective and high-throughput methods for the screen of virus-host interactions.

## Conclusions

This study has developed a Gausian model for predicting prokaryotic virus hosts with better performances than those of VHM and WIsH based on virus genomes. A software tool named Prokaryotic virus Host Predictor was further developed to provide a friendly user interface for the Gaussian model. The work will contribute to the rapid identification of virus hosts in metagenomic studies, and will extend our knowledge of virus-host interactions.

## Abbreviations

GM: Gaussian model
VHM: VirHostMatcher
PHP: Prokaryotic virus Host Predictor
ICTV: International Committee on Taxonomy of Viruses
CRT: CRISPR Recognition Tool.

## Declarations

## Acknowledgments

We thank Prof. Fan Wu in the College of Computer Science and Electronic Engineering in Hunan University and Dr. Jing Meng in the Suzhou Institute of Systems Medicine for the helpful discussions. We thank Ms. Xiarong Yu in the Technische Universität Darmstadt, Ms. Linjie Zhou in the University of Queensland, Ms. Jing Xiong in the Hunan University, Mr. Guanyu Jiang in the University of Minnesota, Mr. Jiarui Xiong in the King’s College London, Mr. Jiakang Xiong in the Starr’s Mill High School, Mr. Qiuhan Jin in the Universität Zürich, Mr. Ruiting Li in the University of Melbourne, Mr. Wu Ding in the Huazhong University of Science and Technology, Mr. Jingbiao Wang in the Tianjin Port and Waterway Engineering Co., Ltd., and Mr. Jing Gao in the Hong Kong University of Science and Technology for the testing of PHP web server.

## Consent for Publication

Not applicable

## Funding

This work was supported by the National Key Plan for Scientific Research and Development of China (2016YFD0500300), Hunan Provincial Natural Science Foundation of China (2018JJ3039, 2019JJ50035, 2020JJ3006), the National Natural Science Foundation of China (31671371 and 81902070) and the Chinese Academy of Medical Sciences (2016-I2M-1-005).

## Availability of Data and Materials

The VHM dataset, the test dataset, and the codes for building, testing, and applications of the GMs in this study are available to the public at GitHub (https://github.com/congyulu-bioinfo/PHP) [22]. The online web server of the PHP is publicly available at http://www.computationalbiology.cn/phageHostPredictor/home.html[31].

## Author’s Contributions

YP designed the study. YP and CL provided the main contribution to the design, implementation, and evaluation of the method, figure preparation, and manuscript text. ZZ, ZC, and ZZZ contributed to the implementation and improvement of the software. ZZ and ZC contributed to the design and implementation of the web server. YQ, AW, TJ, and HZ contributed to the manuscript text. All authors read and approved the final manuscript.

## Competing interests

The authors declare that they have no competing interests.

## Ethics Approval and Consent to Participate

Not applicable

## Notes

### Competing Interest Statement

The authors have declared no competing interest.

