## Additional file for "Prokaryotic virus Host Predictor: a Gaussian model for host prediction of prokaryotic viruses in metagenomics"

### Supplementary Tables

**Table S1**. The host prediction accuracies of GM, VHM and WIsH at taxonomic levels from genus to domain versus the number of top N predictions used when using the consensus method. The highest accuracies among three methods were highlighted in bold.

| | N | 1 | 5 | 10 | 20 | 30 |
|---|---|---|---|---|---|---|
| Genus | WIsH | 0.33 | 0.35 | 0.36 | 0.33 | 0.33 |
| | VHM | 0.31 | 0.36 | 0.38 | 0.39 | 0.39 |
| | GM | **0.34** | **0.39** | **0.42** | **0.44** | **0.45** |
| Family | WIsH | 0.42 | 0.45 | 0.47 | 0.45 | 0.44 |
| | VHM | **0.45** | 0.48 | 0.50 | 0.53 | 0.55 |
| | GM | **0.45** | **0.51** | **0.53** | **0.56** | **0.58** |
| Order | WIsH | 0.45 | 0.53 | 0.56 | 0.55 | 0.53 |
| | VHM | 0.53 | 0.56 | 0.57 | 0.60 | 0.60 |
| | GM | **0.60** | **0.65** | **0.66** | **0.68** | **0.69** |

| | | | | | | |
|---|---|---|---|---|---|---|
| Class | WIsH | 0.69 | 0.72 | 0.75 | 0.77 | 0.76 |
| | VHM | 0.70 | 0.73 | 0.74 | 0.76 | 0.77 |
| | GM | **0.78** | **0.82** | **0.84** | **0.85** | **0.86** |
| Phylum | WIsH | 0.72 | 0.74 | 0.78 | 0.79 | 0.79 |
| | VHM | 0.75 | 0.76 | 0.77 | 0.77 | 0.78 |
| | GM | **0.80** | **0.86** | **0.86** | **0.86** | **0.87** |
| Domain | WIsH | **0.99** | **0.99** | **0.99** | 0.98 | 0.98 |
| | d2 | 0.97 | 0.98 | 0.98 | **0.99** | **0.99** |
| | GM | 0.98 | 0.98 | 0.98 | 0.98 | 0.98 |

**Table S2**. The host prediction accuracies of GM, VHM and WIsH at taxonomic levels from genus to domain *versus* the recall rate when using the threshold method. The highest accuracies among three methods were highlighted in bold.

| Taxonomy level | Recall rate | 0.1 | 0.2 | 0.3 | 0.4 | 0.5 | 0.6 | 0.7 | 0.8 | 0.9 | 1 |
|---|---|---|---|---|---|---|---|---|---|---|---|
| Gneus | WIsH | 0.42 | **0.53** | 0.5 | **0.49** | 0.42 | **0.42** | **0.41** | **0.38** | **0.36** | 0.33 |
| | VHM | **0.58** | 0.43 | 0.38 | 0.38 | 0.38 | 0.36 | 0.35 | 0.34 | 0.32 | 0.31 |
| | GM | 0.52 | **0.53** | **0.51** | 0.46 | **0.45** | **0.44** | **0.41** | **0.38** | **0.36** | **0.34** |
| Family | WIsH | 0.44 | 0.61 | 0.58 | 0.57 | 0.52 | 0.52 | 0.51 | 0.49 | 0.45 | 0.42 |
| | VHM | 0.66 | 0.55 | 0.50 | 0.52 | 0.52 | 0.50 | 0.50 | **0.50** | **0.48** | **0.45** |
| | GM | **0.85** | **0.74** | **0.69** | **0.61** | **0.58** | **0.57** | **0.53** | **0.5** | **0.48** | **0.45** |
| Order | WIsH | 0.45 | 0.62 | 0.59 | 0.58 | 0.54 | 0.54 | 0.53 | 0.51 | 0.49 | 0.45 |
| | VHM | 0.82 | 0.64 | 0.58 | 0.59 | 0.61 | 0.60 | 0.60 | 0.58 | 0.56 | 0.53 |
| | GM | **0.9** | **0.79** | **0.74** | **0.68** | **0.68** | **0.67** | **0.64** | **0.62** | **0.61** | **0.6** |
| Class | WIsH | 0.89 | 0.85 | 0.86 | **0.87** | 0.82 | 0.79 | 0.76 | 0.75 | 0.74 | 0.69 |
| | VHM | **0.98** | **0.90** | **0.87** | 0.86 | 0.85 | 0.81 | 0.80 | 0.78 | 0.74 | 0.70 |

| | | | | | | | | | | | |
|---|---|---|---|---|---|---|---|---|---|---|---|
| | GM | 0.94 | 0.86 | 0.82 | 0.8 | **0.82** | **0.83** | **0.81** | **0.81** | **0.8** | **0.78** |
| | WIsH | 0.90 | **0.93** | **0.93** | **0.93** | **0.88** | 0.84 | 0.81 | 0.78 | 0.77 | 0.72 |
| Phylum | VHM | 0.86 | 0.90 | 0.85 | 0.85 | 0.84 | 0.81 | 0.81 | 0.79 | 0.76 | 0.75 |
| | GM | **0.98** | 0.9 | 0.86 | 0.84 | 0.85 | **0.86** | **0.84** | **0.82** | **0.82** | **0.8** |
| | WIsH | 0.96 | 0.98 | 0.99 | 0.99 | 0.99 | 0.98 | 0.99 | **0.99** | **0.99** | **0.99** |
| Domain | VHM | **1.00** | **1.00** | **1.00** | **1.00** | **1.00** | **1.00** | **1.00** | 0.98 | 0.98 | 0.97 |
| | GM | **1.00** | 0.97 | 0.98 | 0.98 | 0.98 | 0.99 | 0.99 | **0.99** | 0.98 | 0.98 |

**Table S3**. Comparison of the time consumed in building models and prediction of virus hosts of PHP and WIsH on the VHM dataset.

**Part I**. The comparison was conducted on a laptop with the Operating System of Ubuntu 16.04 LTS, with CPU of Intel i5 (9300H, 2.40GHz, 4 cores and 8 threads), with a RAM of 24 GB, and with a hard disk of SSD.

| Method | Time consumed in the model building | Time consumed in prediction of virus hosts | Total time consumed |
|---|---|---|---|
| PHP | 3h 30m | 57m | 4h 27m |
| WIsH | 27m | 1h 51m | 2h 18m |

**Part II**. The comparison was conducted on a server with the Operating System of Ubuntu 16.04 LTS, with CPU of Intel Xeon (Silver 4114, 2.20GHz, 20 cores and 40 threads), with a RAM of 128GB, and with a hard disk of SSD.

| Method | Time consumed in the model building | Time consumed in prediction of virus hosts | Total time consumed |
|---|---|---|---|
| PHP | 48m | 13m | 1h 1m |
| WIsH | 12m | 1h 2m | 1h 14m |

**Supplementary Figures**

**Figure S1.** The comparisons of the taxonomic distribution of both viruses and hosts in the VHM and test datasets used in the manuscript

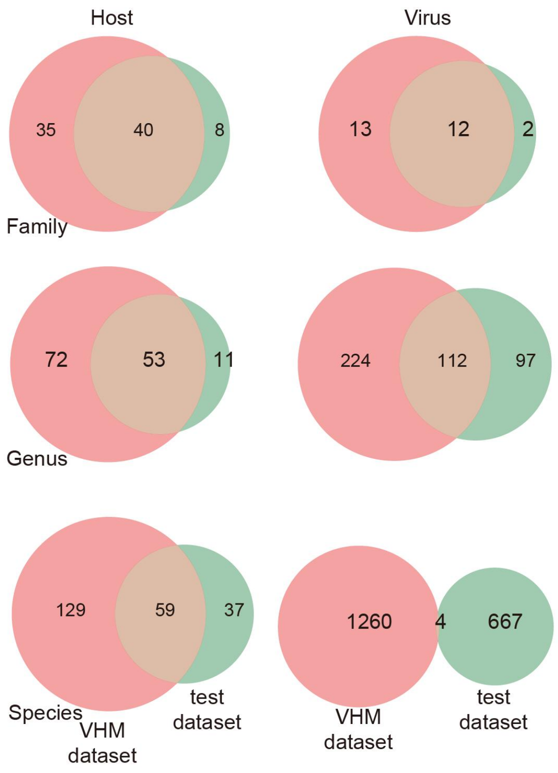


**Figure S2.** The host prediction accuracy of the Gaussian mixture model in the ten-fold cross-validations on the K-means clustering of the VHM dataset *versus* the length of K-mer (A) and the number of components in the Gaussian mixture model(B).

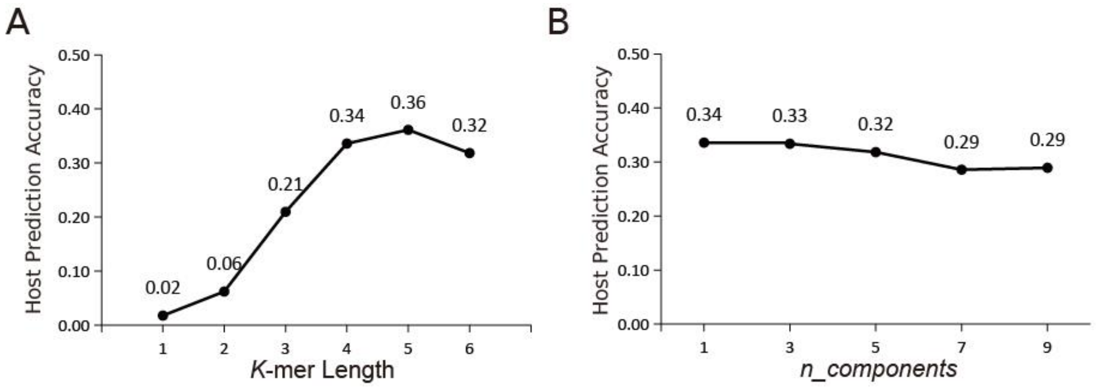


**Figure S3**. The comparison between GM and other common machine learning algorithms on host prediction accuracy. All models took the *k*-mer frequency (k=4) as features, and were trained with the default parameters. Negative samples are needed for building computational models with these machine-learning algorithms, and were obtained as follows: for each virus, a non-host prokaryote was randomly selected, which resulted in the same number of negative samples as the positive samples, i.e, virus-host interactions. The testing strategy mentioned in Figure 1 was used to evaluate the prediction ability of these machine-learning algorithms. During the testing process, only the positive samples, i.e., virus-host interactions, were clustered since we aimed to predict viral hosts. The random forest (RF) algorithm was selected for further optimization since it was observed to perform best among these machine-learning algorithms. Two parameters, i.e., the number of decision trees (*n_estimators*) and the number of features to consider when looking for the best split (*max_features*), have a key impact on the performance of the RF algorithm. Besides, the length of *k*-mers used in the modeling may also have an influence on the performance of the RF algorithm. Therefore, the *n_estimators*, *max_features*, and the length of *k*-mers were further tuned to improve the RF algorithm. The RF algorithm with the "n_estimators" set to be 2000, "max_features" set to be "auto", and *k*-mer length set to be 6 was found to perform best (see "Random forest with best parameters" in the figure). The modeling with these machine-learning algorithms was computed with the "scikit-learn" package in python.

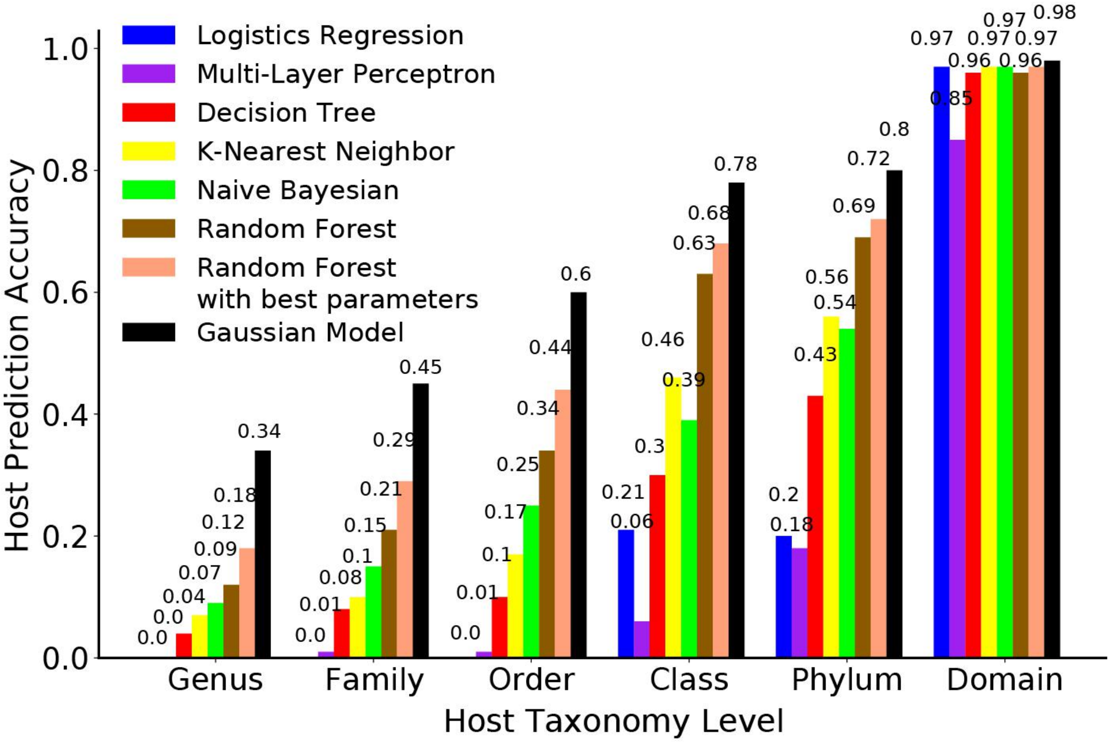


**Figure S4**. The distribution of k-mer frequency differences between virus and hosts genomes (blue), and those between virus and non-host prokaryotic genomes (orange). (A-H) represents the randomly selected k-mers of AGTT, AAAA,TTGC, CACG, AATT, TAGA, CGGG, TTCT, respectively.

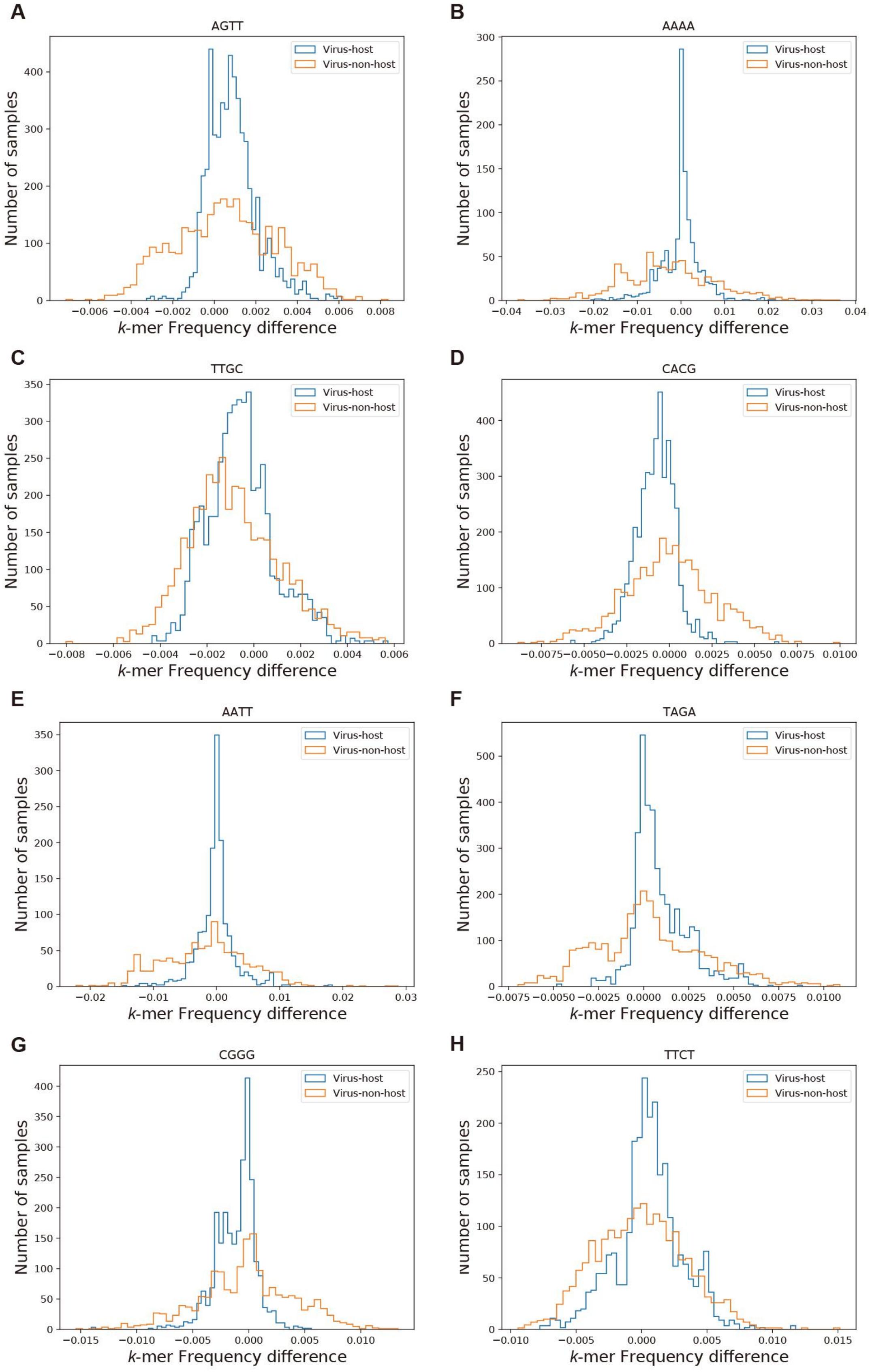
A
AGTT
B
AAAA
C
TTGC
D
CACG
E
AATT
F
TAGA
G
CGGG
H
TTCT
Virus-host
Virus-non-host
Number of samples
k-mer Frequency difference

**Figure S5**. The taxonomic distribution of viruses in the VHM dataset.

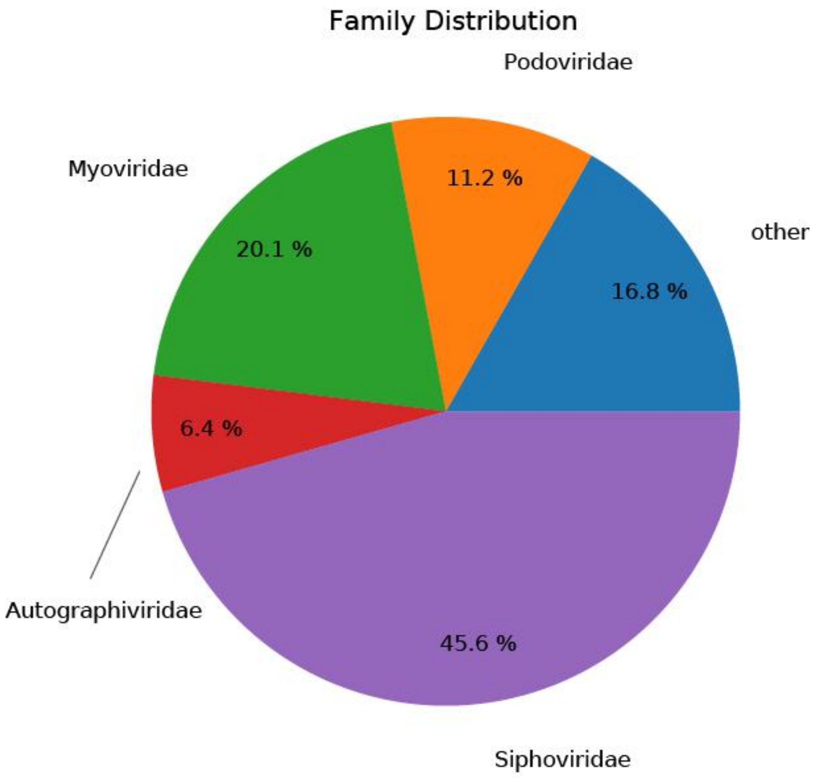
